# Frontal eye field inactivation reduces saccade preparation in the superior colliculus, but does not alter how preparatory activity relates to saccade latency

**DOI:** 10.1101/215137

**Authors:** Suryadeep Dash, Tyler R. Peel, Stephen G. Lomber, Brian D. Corneil

## Abstract

A neural correlate for saccadic reaction times (SRTs) in the gap saccade task is the level of preparatory activity in the intermediate layers of the superior colliculus (iSC) just before visual target onset: greater levels of iSC preparatory activity precede shorter SRTs. The frontal eye fields (FEF) are one likely source of such iSC preparatory activity, since FEF preparatory activity is also inversely related to SRT. To better understand the FEF’s role in saccade preparation, and the way in which such preparation relates to SRT, in two male rhesus monkeys we examined iSC preparatory activity during unilateral reversible cryogenic inactivation of the FEF. FEF inactivation increased contralesional SRTs, and lowered ipsilesional iSC preparatory activity. FEF inactivation also reduced fixation-related activity in the rostral iSC. Importantly, the distributions of SRTs generated with or without FEF inactivation overlapped, enabling us to conduct a novel population-level analyses examining iSC preparatory activity just before generation of SRT-matched saccades. These analyses revealed no change during FEF inactivation in the relationship between iSC preparatory activity and SRT-matched saccades across a range of SRTs, even for the occasional express saccade. Thus, while our results emphasize that the FEF has an overall excitatory influence on preparatory activity in the iSC, the communication between the iSC and downstream oculomotor brainstem is unaltered for SRT-matched saccades, suggesting that the integration of preparatory and visual signals in the SC just before saccade initiation is largely independent of the FEF for saccades generated in this task.

**Significance statement:** How does the brain decide when to move? Here, we investigate the role of two oculomotor structures, the superior colliculus (SC) and frontal eye fields (FEF), in dictating visually-guided saccadic reaction times (SRTs). In both structures, higher levels of preparatory activity precede shorter SRTs. Here, we show that FEF inactivation increases SRTs and decreases SC preparatory activity. Surprisingly, a population-level analysis of SC preparatory activity showed a negligible impact of FEF inactivation, providing one examines SRT-matched saccades. Thus, while the FEF is one source of preparatory input to the SC, it is not a critical source, and it is not involved in the integration of preparatory activity and visual signals that precedes saccade initiation in simple visually-guided saccade tasks.

## Introduction

The time to respond to a behavioral event can be highly variable. Even for simple visually-guided saccades, saccadic reaction times (SRTs) can range from those approaching the minimal sensory-to-motor delays in the case of express saccades to others that are two to three times longer (Saslow 1967; Fischer et al. 1984; Paré and Munoz 1996). One neural correlate of SRT variability in this task is the level of low-frequency preparatory activity in the intermediate layers of the caudal superior colliculus (iSC) attained just before the arrival of visual information arising from target presentation. Across a variety of tasks which manipulate top-down signals, systematically greater levels of preparatory activity precede progressively shorter SRTs (Dorris et al. 1997; Basso and Wurtz 1998; Krauzlis 2003; Rezvani and Corneil 2008; Marino et al. 2015). Importantly, the level of preparatory activity does not relate to variability in other saccadic parameters like saccade endpoint or peak velocity (Basso and Wurtz 1998).

The goal of the current study is to investigate how the relationship between preparatory iSC activity and SRT is altered when the frontal eye fields (FEF), a source of top-down information to the iSC (Everling and Munoz 2000; Sommer and Wurtz 2000, 2001; Wurtz et al. 2001), are temporarily inactivated. Doing so will allow us to address two questions. First, by combining FEF inactivation with iSC recordings, we can address the importance of FEF integrity on iSC preparatory activity. Second, and more importantly, this approach will allow us to examine the relationship between iSC preparatory activity and SRT when the oculomotor system is in an altered state. FEF inactivation increases SRTs, but how such increased SRTs relate to preparatory iSC activity is unknown. Are SRTs increased simply because FEF inactivation decreases preparatory activity? If so, then the relationship between iSC preparatory activity and SRT would simply shift to higher SRTs, so that preparatory activity would remain the same for saccades of matched SRTs. Alternatively, perhaps more preparatory activity is required to generate a saccade of a given SRT, perhaps due to the loss of FEF signaling along parallel circuits to the oculomotor brainstem that bypass the iSC (Raybourn and Keller 1977; Leichnetz 1980, 1981; Segraves 1992), or to an increase in activity in the rostral iSC, given the reciprocal relationship between fixation-related and preparatory activity in the rostral and caudal iSC, respectively (Dorris and Munoz 1995; Dorris et al. 1997; Munoz and Istvan 1998; Munoz and Fecteau 2002; Takahashi et al. 2010).

We combined unilateral cryogenic inactivation of the FEF with recording of preparatory-activity in the ipsilesional caudal iSC and fixation-related activity in the rostral iSC while monkeys performed a visually-guided gap saccade task. Cryogenic inactivation allowed us to follow the changes in iSC activity immediately (within minutes) following FEF inactivation, hence any changes in iSC activity are unconfounded by functional recovery that occurs in the days, weeks, or months after permanent lesions of the FEF (Schiller et al. 1987). Our choice of a visually-guided saccade task was advantageous: although the changes in saccadic behavior are less than those observed for delayed saccades (Peel et al. 2014), the distributions of SRTs made with or without FEF inactivation overlap considerably. This allows us to investigate how low-frequency preparatory iSC activity relates to SRT with and without FEF inactivation, both within single neurons and across the recorded population for SRT-matched saccades. We found that FEF inactivation increased SRTs, and decreased both preparatory activity in the caudal iSC and fixation-related activity in the rostral iSC. However, we found that the level of iSC preparatory activity did not change when we examined the subset of SRT-matched saccades. These results show that the relationship between iSC preparatory activity and SRT is unaltered during FEF inactivation, so that a saccade of the same SRT, including those generated at express-saccade latencies, can be generated providing other non-FEF sources compensate for the loss FEF signaling.

## Methods

### Subjects and surgical procedures

Two male rhesus monkeys (*Macaca mulatta*, DZ, and OZ weighing 9.8, and 8.6 kg respectively) were prepared for head immobilization, cryogenic inactivation of FEF and electrophysiological recordings from the iSC. All training, surgical, and experimental procedures were in accordance with the Canadian Council on Animal Care policy on the use of laboratory animals (Olfert et al. 1993) and approved by the Animal Use

Subcommittee of the University of Western Ontario Council on Animal Care. We monitored the monkeys’ weights daily and their health was under the close supervision of the university veterinarians. Surgical procedures describing drug regimes, post-surgical care implantation of head post, cryoloops, and placement of recording chamber have been described previously (Rezvani and Corneil 2008; Peel et al. 2014). Details of the cryoloop dimensions, placement method and volume of inactivation for the monkeys used in this study can be found elsewhere (Peel et al. 2014, 2016, 2017). Briefly, each monkey was implanted bilaterally with two stainless steel cryoloops in the inferior and superior aspects of the arcuate sulcus [inferior arm (IA), superior arm (SA)],). In this study we have only used unilateral IA cooling in both monkeys as this increases trial yield during cooling, and produces ~70% of the SRT deficits caused by combined unilateral cooling of the IA and SA (Peel et al. 2014).

### Experimental procedures

Monkeys were seated in a custom-made primate chair with their head immobilized and faced a rectilinear grid of >500 red LEDs covering ± 35˚ of the horizontal and vertical visual field. The eye movements were recorded using a single, chair-mounted eye tracker (EyeLink II, resolution = 0.05˚, sampling rate 500Hz). All experiments were conducted in a dark, sound-attenuated room. The behavioral tasks were controlled by customized real-time LabView programs on a PXI controller (National Instruments) at a rate of 1 kHz. Extracellular single-unit activity was recorded with epoxylite insulated tungsten microelectrodes (0.5–2 MΩ at 1 kHz; FHC, Bowdoin, ME) lowered through 23-gauge guide tubes and advanced to the dorsal surface of the SC using a microdrive (NAN instruments). Neural activity was amplified, filtered, and stored for off-line sorting via a Plexon MAP system (Plexon, Dallas, TX).

An experimental dataset consisted of pre-cooling, peri-cooling, and post-cooling sessions, with each session containing 60-120 correct trials (requiring between 180-360 trials total). After the pre-cooling session, the cooling pumps were turned on allowing the flow of chilled methanol through the lumen of the cryoloops. The peri-cooling session was initiated when cryoloop temperature reached and stayed stable at 3°C.

Once sufficient data was collected for the peri-cooling session, cooling pumps were turned off, which allowed the cryoloop temperature to rapidly return towards body temperature. When cryoloop temperature exceeded 35˚C, the post-cooling session was initiated. To control for time dependent factors like satiation, we pooled data from pre-cooling and post-cooling session and termed this the *FEFwarm* condition. Data collected during FEF inactivation was termed as coming from the *FEFcool* condition.

### Behavioral tasks

Monkeys performed intermixed visually-guided gap and no-gap saccades to peripheral targets placed either in the center of the neuron’s response field (RF) or in the diametrically opposite location. As this study concerns saccade preparatory activity in caudal iSC and fixation activity in rostral iSC, we only present data from the gap saccade task. Monkeys fixated the central fixation LED for 750-1000ms until it disappeared. On gap trials, after a blank period of 200ms (gap interval), during which the monkeys maintained central fixation, a peripheral target appeared for 150ms to which the monkeys had to saccade as quickly as possible. No-gap trials were the same but without a 200ms gap interval, i.e., fixation LED disappearance coincided with peripheral target’s appearance. Saccades made within a temporal window of 500ms after peripheral target appearance within a circular spatial window (diameter 60% of target’s visual eccentricity) received fluid reward. The large tolerance window was based on the expected spatial deficits that follow FEF inactivation (Sommer and Tehovnik 1997; Dias and Segraves 1999; Peel et al. 2014). However, subsequent analysis (see results) revealed only small changes in end-point saccadic scatter during FEF inactivation.

A recent study suggested that low frequency activity in the caudal iSC during a delayed saccade task may carry additional information about saccade kinematics in addition to preparatory signals (Jagadisan and Gandhi 2017). There is an important distinction between our study and that of Jagadisan and Gandhi as the knowledge of the saccade goal precedes the buildup of low frequency activity in their task. Moreover, a previous study employing an immediate response task similar to ours did not find any correlation between preparatory activity with saccade metric or kinematic (Basso and Wurtz 1998). Also, we (in this study) and others have also shown that the overall level of preparatory activity for ipsi- and contraversive saccades is not different when their probability of occurrence is same (Dorris et al. 1997; Sparks et al. 2000; Krauzlis 2003; Rezvani and Corneil 2008).

All behavioral analyses were carried out using customized MATLAB programs (MATLAB, The MathsWorks Inc., MA). Eye position traces were filtered using a 3rd order low pass butterworth filter and differentiated to produce eye velocity. Eye velocity was used to determine the onset and offset times of saccades with a velocity criterion of 30°/s and the maximum instantaneous velocity between saccade onset and offset was deemed as peak velocity. The eye position at saccade offset was used to calculate the mean saccade amplitude (horizontal and vertical) and end-point scatter which represented the mean angular distance between the displacements of mean and individual saccade end points from the central fixation position (White et al. 1994). We analyzed the first saccade following fixation LED disappearance. Visual inspection of the data off-line confirmed the validity of automatic marking. Trials with SRT <50ms were classified as anticipatory and discarded.

### Neuron classification

We estimated the instantaneous firing rate of a recorded neuron with a continuous spike density function for each trial, generated by convolving the spike train with a postsynaptic activation function with a rise time of 1ms and a decay time of 20ms (Hanes et al. 1995). Use of a different convolution function (e.g., a 10ms gaussian kernel) did not alter any of the results. We classified our sample of neurons broadly based on a previous study (Dorris et al. 1997). Caudal iSC neurons were classified as saccade-related if they had a peak firing rate >100 spikes/s around saccades (-20ms to 20ms relative to saccade onset) made into the RF. Saccade related neurons were further classified as build-up neurons or burst neurons based on the presence or absence, respectively, of preparatory activity (PREP) attained just before the arrival of visual information. For a given saccade related neuron, the average activity was computed in three different time epochs: *baseline* activity was sampled between - 500ms to −200ms before target onset; *PREP* activity was sampled between −70 to +30ms relative to target onset, and the final level of PREP activity (Final PREP) was sampled in the last 20ms of PREP epoch. In our sample of neurons, the visual transient did not arrive until at least 40ms after target onset in any of the neurons based on Poisson burst onset detection method described elsewhere (mean visual onset time across neurons= 45ms± 3ms; minimum average visual onset time=40ms; Hanes et al. 1995; Peel et al. 2017). A neuron was deemed to exhibit significant PREP activity if the average PREP activity was higher than average baseline activity during the gap-saccade task (p< 0.05; Wilcoxon sign rank test; pooled across both target directions). We also recorded neurons from the rostral pole of iSC. For rostral iSC fixation neurons, we calculated the same parameters as the buildup neurons but named it differently: *baseline* activity sampled between −500ms to −200ms before target onset; *FIX* activity sampled between −70 to +30ms relative to target onset and final level of FIX activity (Final FIX) sampled in the last 20ms of FIX epoch. We defined rostral iSC neurons as fixation neurons if they fulfilled the criterion of exhibiting tonic average firing rate of >10spikes/s during both baseline and FIX epoch during gap saccade task along with a significant decrease in activity at saccade onset (Dorris et al. 1997). Although rostral pole iSC neurons also encode microsaccades (Hafed et al. 2009; Krauzlis et al. 2017) very few microsaccades are generated during the time intervals of interest in this task (Watanabe et al. 2014).

### PREP-SRT correlation

We also performed trial-by-trial linear regression (Pearson’s correlation) between final PREP and contralateral SRT in buildup neurons and compared the coefficient of determination (r^2^), y-intercept and slope of this relationship across FEF inactivation. A linear regression between PREP and SRT yielded qualitatively similar results. For the rostral iSC fixation neurons, a trial by trial linear regression between either final FIX or FIX with SRT did not yield significant correlations.

### Detection of onset of preparatory activity

Recently, we reported that change in SRT following FEF inactivation during a delayed saccade task related best to changes in the onset time of saccadic accumulation (Peel et al. 2017). In this study we also investigated if there were changes in the onset time of PREP accumulation. To detect the onset time of PREP accumulation on a neuron-by-neuron basis, we implemented a piecewise two-piece linear regression of PREP iSC activity using an approach similar to Peel and colleagues (2017). The objective of this analysis was to find the two linear regressions that best fit the convolved average iSC PREP activity before the onset of the visually-related activity following target onset. The first linear regression is based on activity from 200ms before the gap onset to a candidate inflection point; the second linear regression is based on activity from this candidate inflection point to the peak of PREP activity during the final PREP epoch (10- 30ms after visual target onset). For rostral iSC fixation neurons exhibiting a decrease in activity during the gap interval, the second linear regression ran from the candidate inflection point to the trough of FIX activity. Candidate inflection points were tested every millisecond from 200 ms before gap onset to 30 ms after target onset. The onset of preparatory activity was taken as the inflection point that minimized the summed squared error between convolved iSC activity and the two linear regressions.

### Experimental design and statistical analysis

In this study, we compare both behavior (SRT, peak velocity, saccade amplitude and end-point scatter of saccades) and neuronal activity across FEF inactivation. Single session comparisons (FEF*warm* vs FEF*cool*) of either behavior or single neuron activity were performed using a Wilcoxon rank sum test at a p level of 0.05. For buildup neurons, we compared activity during the baseline, PREP, final PREP intervals, onset time of PREP accumulation and compared the slope of PREP, which was calculated by differentiating the spiking activity in the PREP interval. For the rostral iSC, corresponding time epochs, slope and onset times of the FIX activity was compared. To assess changes in SRT and neural activity across the sample of recorded iSC neurons following FEF inactivation, we performed paired group level comparisons using a Wilcoxon sign rank test at a p level of 0.05. We also implemented a novel population-based analysis comparing population PREP activity for SRT matched saccades across FEF inactivation. The logic and description of this analysis is described in the Results.

## Results

We report single unit activity from the left iSC of 2 monkeys (DZ and OZ) while the left FEF was reversibly inactivated by cryogenic means, when animals were engaged in the gap saccade task. We performed 54 inactivation sessions in monkey DZ and 35 inactivation sessions in monkey OZ. In each session for a neuron to be further analyzed, between 15-30 valid trials per target location in each of the pre-, peri- and post-cooling sessions was deemed necessary. Across all the inactivation sessions (n=89 sessions), 48 caudal iSC saccade-related neurons (27 from monkey DZ and 21 from monkey OZ) could be isolated for long enough to collect sufficient pre-, peri-, and post-cooling data. Of these 48 neurons, 35 exhibited significant preparatory activity (PREP), and were classified as buildup neurons (18 and 17 neurons from monkey DZ and monkey OZ, respectively). Response fields centers for the buildup neurons ranged between 4°/6° to 20°/20° in horizontal/vertical coordinates relative to fixation point with the mean response center of 9° ± 3.7° (horizontal, mean ± sd) and 2.5° ± 5.8° (vertical). Fixation-related activity was also recorded from 17 rostral iSC neurons (9 and 8 neurons from monkey DZ and OZ, respectively).

Consistent with our previous reports (Peel et al. 2014, 2016, 2017), unilateral cryogenic FEF inactivation increased contraversive SRTs within and across all sessions (Fig. 1B, D). In contrast to previous reports, such inactivation did not consistently increase ipsiversive SRTs (Fig. 1A, C). The failure to observe ipsiversive SRT increases may be due to differences in target configuration and behavioral task (e.g., 2 potential targets here versus 32 potential targets in the visually-guided saccades tasks reported in (Peel et al. 2014); the use of a visually-guided saccade task here versus the use of delayed saccade tasks in (Peel et al. 2014, 2016, 2017)). We also compared saccade metrics and kinematics across all the sessions. We did not observe any change in horizontal saccade amplitude across FEF inactivation for all sessions (n=89 sessions, mean difference in horizontal amplitude=0.07°±0.61°; p=0.3964, z-val=0.8480; Sign test). Vertical saccade amplitude showed a significant, albeit small, decrease (mean difference in vertical amplitude=-0.37°±0.82°; p=0.0015, z-val=-3,18; Sign test). The mean endpoint scatter of saccades did not change in 59 out of 89 sessions (p>0.05; Wilcoxon rank sum test) and across all sessions also showed only a small but significant increase from 1.04±0.30° to 1.27±0.34° (p=2.42e-08, z-val=-5.5785; Wilcoxon sign rank test). Next, we compared peak saccade velocity across session and found a reduction in peak saccade velocity in 63 out of 89 sessions during FEF inactivation (p<0.05; Wilcoxon rank sum test). Across sessions, mean saccade peak velocity decreased significantly (14% decrease, p=3.32e-15, z-val=7.8780; Wilcoxon sign rank test).

**Figure 1:**
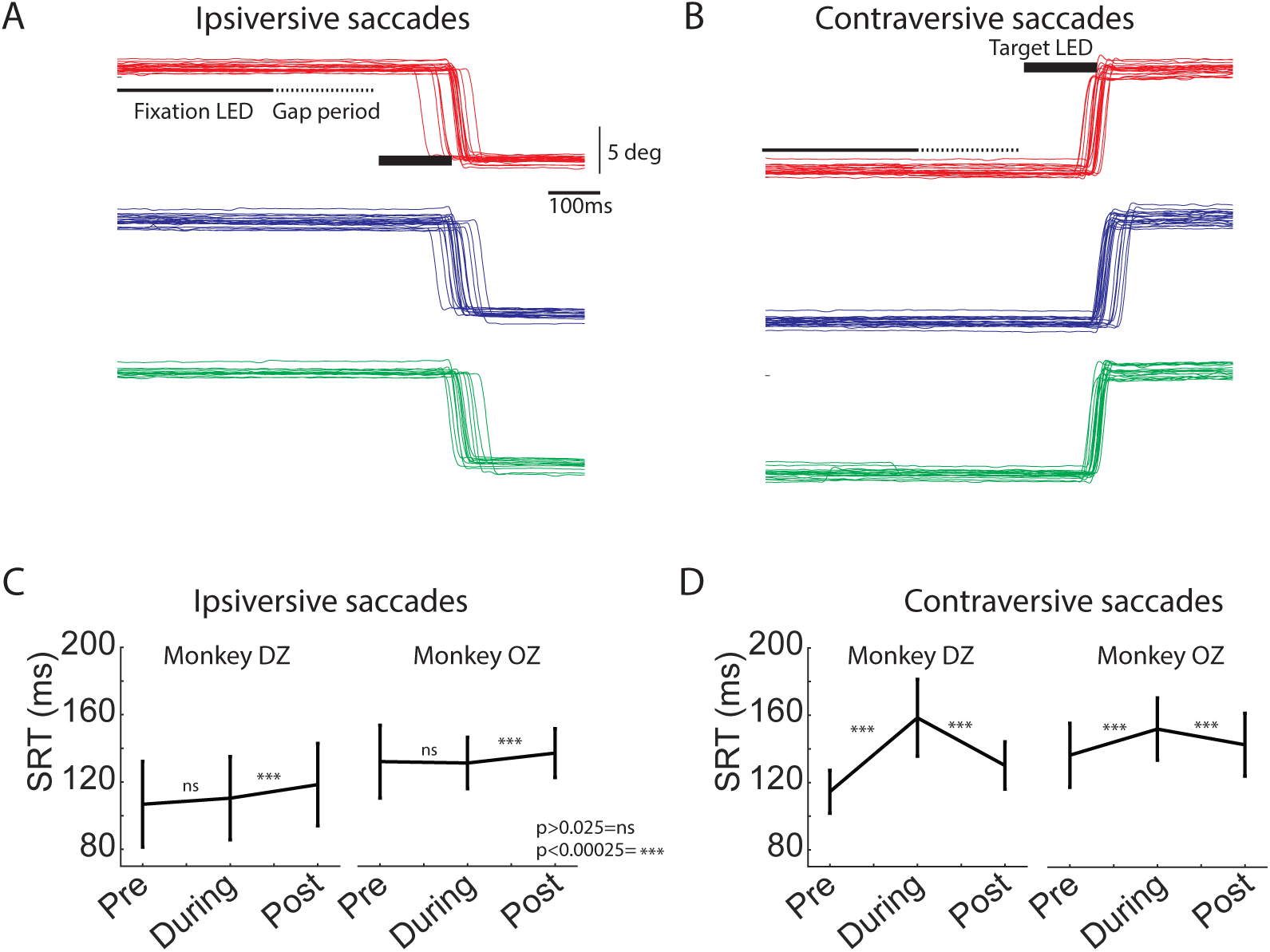
FEF inactivation increased SRTs for contraversive saccades. A & B: Eye position traces for an example session across pre- (red), during (blue) and post (green) cooling trials for ipsiversive (A: leftward saccades) and contraversive (B: rightward saccades) saccades, respectively (left FEF was inactivated). C & D: Each line connects mean SRT (± SD) across pre-, during, and post cooling sessions for each of the 2 monkeys for ipsiversive (C) and contraversive (D) saccades. Both monkeys exhibited increased SRT for contraversive saccades during FEF inactivation that rebounded after FEF rewarming.

### FEF inactivation reduced preparatory activity in ipsilesional iSC

We next investigated how the increase in SRT during FEF inactivation related to changes in iSC preparatory (PREP) activity. Such activity precedes the arrival of visual information into the iSC, and hence is driven strictly by top-down factors such as the expectation of target appearance and/or bottom-up factors like fixation disengagement due to the 200 ms gap (Reuter-Lorenz et al. 1991; Munoz and Wurtz 1992; Tam and Ono 1994; Dorris and Munoz 1995; Paré and Munoz 1996). Figures 2A-C show activity aligned to contralateral target presentation for three example caudal iSC buildup neurons, to emphasize the diversity of effects accompanying FEF inactivation. Contraversive SRTs increased significantly during FEF inactivation in all three examples (p=4.62e-05, z-val=-4.737; p=1.14e-07, z-val=-5.3022 and p=0.01, z-val=-2.55, respectively; Wilcoxon rank sum test; horizontal boxplots in Figs. 2A-C). Note how the activity of these neuron started to rise ~100ms before target onset (or ~100ms after FP offset), and then sharply increased ~40ms after target presentation due to the arrival of visual information from the contraversive target. The activity of all caudal iSC neurons sharply decreased ~40 ms after ipsilateral target presentation (results not shown). In this manuscript, we are primarily interested in the initial rise of iSC activity during the gap period, prior to the arrival of visual information (zoomed interval in Figs. 2A-C). As expected, all aspects of activity within this interval (e.g., activity during baseline, PREP, final PREP and rate of accumulation of PREP activity) did not differ significantly on trials with ipsi-versus contraversive targets (p < 0.05 for all comparisons, Wilcoxon rank sum test).

**Figure 2:**
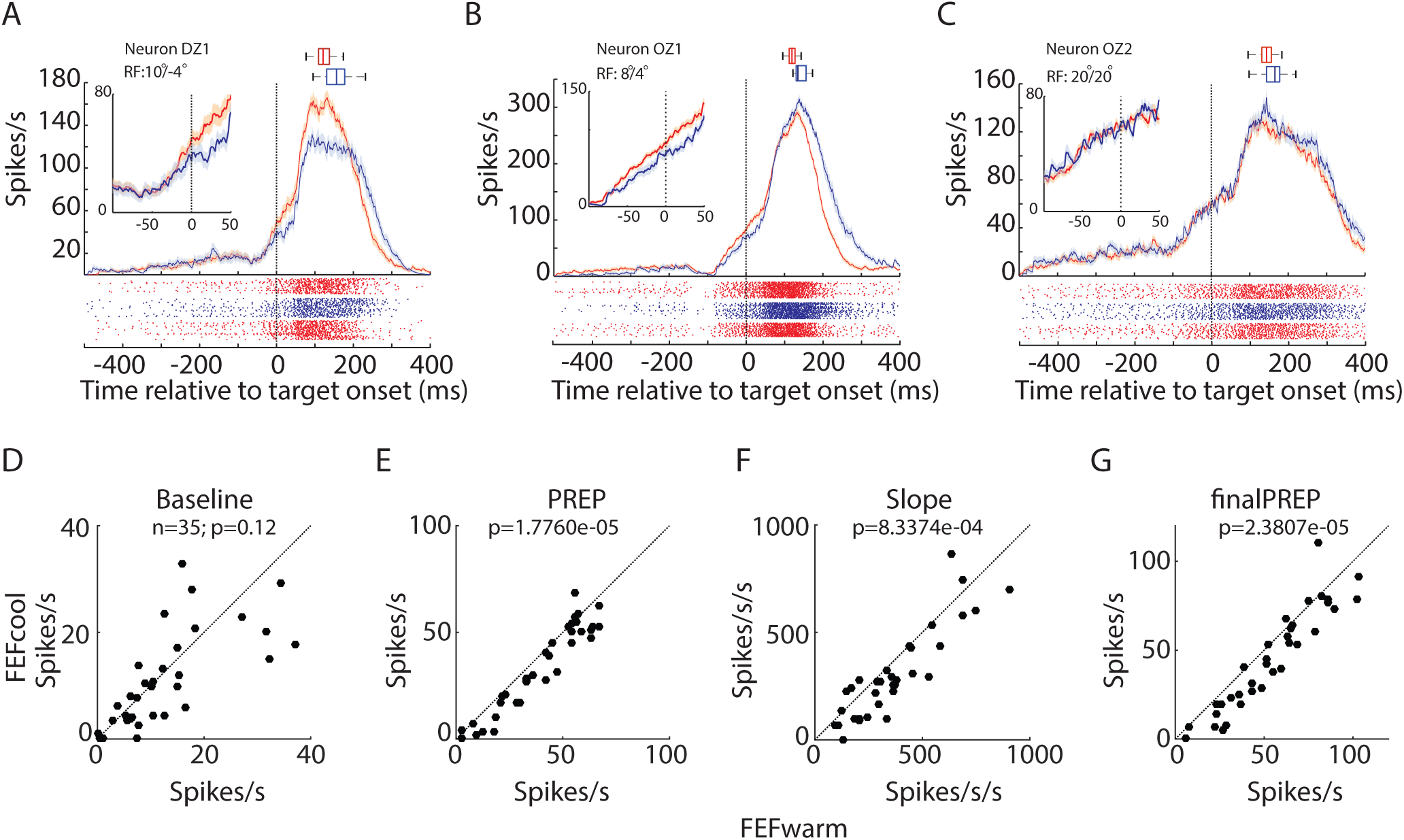
FEF inactivation decreased preparatory activity in the ipsilesional iSC. (A, B & C) Spike rasters (below) and mean spike density functions (above, mean ± SE) showing reduced (A & B) or unchanged preparatory activity (C) during FEF inactivation. Inset shows the magnified view of the relevant period of interest. Box plots shows the SRT changes for the corresponding behavioral sessions (median, 25-75 percentile and the range). All the sessions showed significant increases in SRT (p<0.05, Wilcoxon rank sum test). Pre- and post-cooling trials pooled as FEFwarm (red) and trials during FEF inactivation are shown as FEFcool (blue). Across all the neurons (n=35), FEF inactivation did not alter baseline activity (D), but reduced PREP activity (E), the slope of PREP activity (F), and the final level of PREP activity (G) at p level of 0.05 (Wilcoxon sign rank test).

For the neurons shown in Figs. 2A & B, FEF inactivation significantly reduced the final level of the PREP from 69 spikes/s and 116 spikes/s during the FEF*warm* condition to 47 spikes/s and 94 spikes/s during the FEF*cool* condition, respectively (p=0.004, z-val=2.8593 and p=0.003, z-val=2.9758, respectively; Wilcoxon rank sum test). The slope of the accumulation of PREP also decreased significantly for these neurons (p=0.0017, z-val=3.1405 and p =0.02, z = 2.9758, respectively; Wilcoxon rank sum test) although baseline activity did not change (p=0.19, z-val=-1.3055 and p=0.4773, z-val=0.7107, respectively; Wilcoxon rank sum test). In contrast, for the neuron shown in Fig. 2C, none of the baseline (p=0.27, z-val=-1.1005), PREP activity (p=0.98, z-val=- 0.0155), final PREP (p=0.41, z-val=0.8214), or slope of accumulation of PREP (p=0.33, z-val=0.9764) changed significantly during FEF inactivation (Wilcoxon rank sum test) despite a significant increase in contraversive SRT during this session (141ms to 160ms, p=0.01, z-val=-2.55; Wilcoxon rank sum test). Across our sample of 35 neurons, at the single neuron level, 9 (25%), 14 (40%), 14 (40%) and 6 (17%) neurons exhibited significant decreases in baseline, PREP, final PREP and slope of PREP, respectively (p < 0.05, Wilcoxon rank sum test).

Next, we analyzed the changes in PREP activity in caudal iSC neurons at the group level. Across our sample of 35 neurons, FEF inactivation did not significantly alter baseline activity (Fig. 2D, p = 0.12, z-val=1.5396; Wilcoxon sign rank test), but lowered overall PREP activity (Fig. 2E, p =1.7e-005, z-val=4.2913), the rate of accumulation of PREP activity (Fig. 2F, p =8.3e-004, z-val=3.3413) and the final PREP attained just prior to the arrival of visual information (Fig. 2G; p =2.3e-005, z-val=4.2258). Thus, FEF inactivation produced a widespread decrease in PREP activity in the caudal iSC.

In a recent study, we showed that FEF inactivation delayed the onset of saccadic activity in a delayed saccade task (Peel et al. 2017). We wondered if the onset of PREP activity during the gap interval would also be modified during FEF inactivation. There did not appear to be any obvious changes in the onset of PREP activity in the representative neurons shown in Fig. 2A-C. To quantify this across our sample, we performed a two-piece linear regression analysis to detect the onset of PREP accumulation (see Methods) and found no change in the onset time of PREP activity across our sample of neurons (n=35; p=0.81; z-val=-0.4472; Wilcoxon sign rank test).

### FEF inactivation reduced fixational activity in ipsilesional rostral iSC

The rostral and caudal portions of the iSC are thought to be mutually antagonistic (Munoz and Istvan 1998; Munoz and Fecteau 2002; Takahashi et al. 2010). Given this, could the observed decreases in PREP activity in the caudal iSC during FEF inactivation be related to increases in fixation-related activity in the rostral iSC? We recorded from 17 rostral iSC neurons that met the classification of fixation-related neurons (9 and 8 neurons from monkey DZ and monkey OZ, respectively), and compared baseline, FIX, slope of FIX activity and the final level of FIX activity across FEF inactivation. Figure 3A-C shows three examples of rostral iSC fixation-related neurons that were highly active during the baseline interval, but then became both less active during the gap interval and silent around the time of a saccade. For all three neurons, FEF inactivation decreased the level of baseline activity (Figs 3A-C). A further reduction in FIX activity was observed in the neurons shown in Figs. 3A and 3B, but not in the neuron shown in Fig. 3C. Further, FEF inactivation decreased the post-saccadic activity on the neuron shown in Fig. 3C, even though the saccade target was extinguished by the time the saccade was generated.

**Figure 3:**
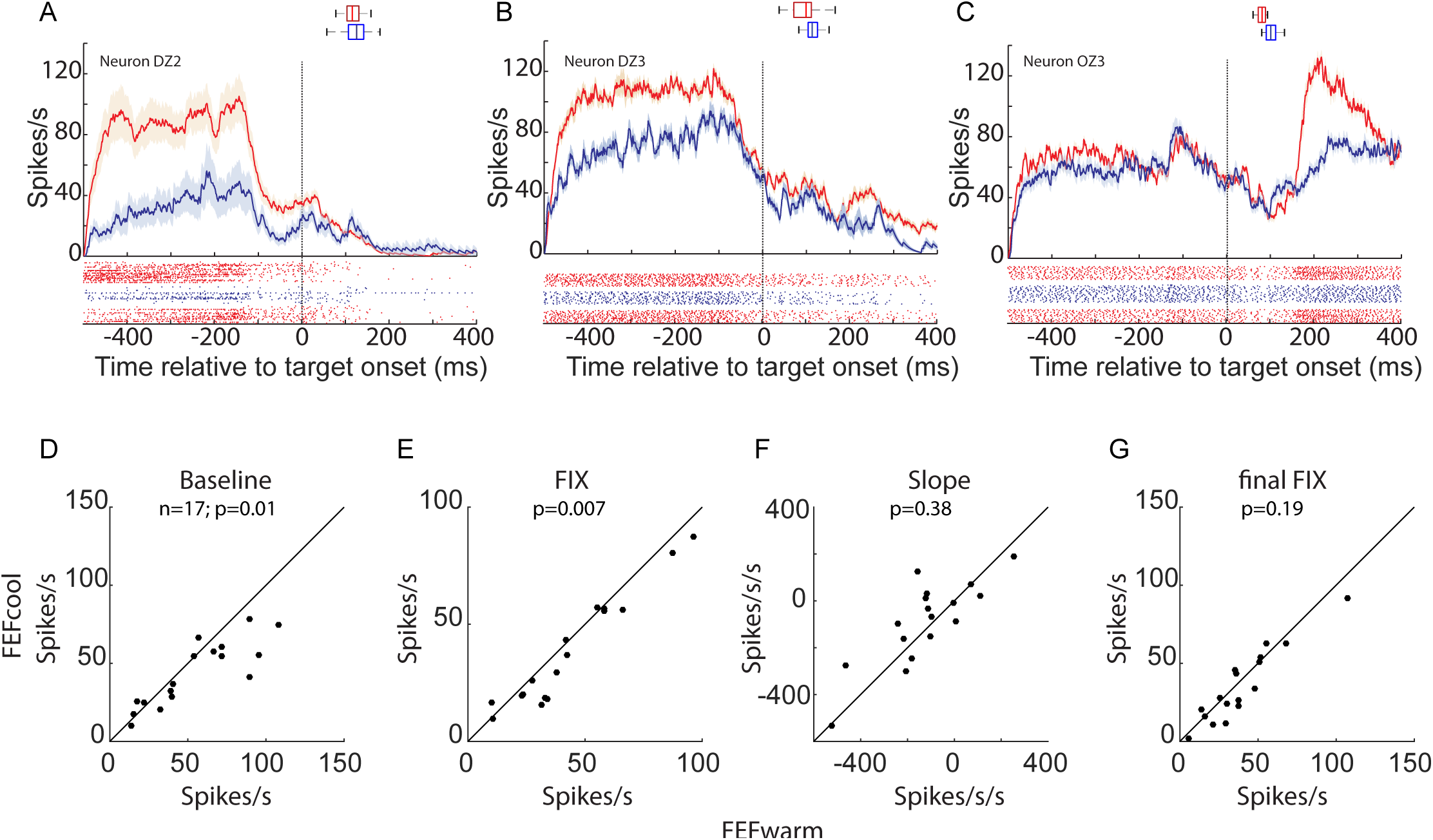
FEF inactivation decreased fixational activity in the ipsilesional rostral iSC. (A, B & C) Spike rasters (below) and mean spike density functions (above) showing reduced fixational baseline activity for three representative neurons. Across all the neurons (n=17), FEF inactivation reduced baseline activity (D), and FIX activity (E), but did not alter the slope of FIX activity (F) or the final level of FIX activity (G) at p level of 0.05 (Wilcoxon sign rank test). Same format as Fig. 2

Next, we analyzed the changes in fixation activity in rostral iSC neurons at the group level. Across our sample of 17 neurons, FEF inactivation significantly altered baseline (Fig. 3D, p = 0.0099, z-val=2.5799; Wilcoxon sign rank test), and FIX activity (Fig. 2E, p =0.007, z-val=2.6746). However, the rate of accumulation of FIX activity (Fig. 2F, p =0.38, z-val=-0.8758) and the final FIX attained just prior to the arrival of visual information (Fig. 2G; p =0.19, z-val=1.3018) were not changed due to FEF inactivation. Thus, in contrast to what would have been expected based on a reciprocal relationship between the caudal and rostral iSC, FEF inactivation reduced, rather than increased, fixation-related activity in the rostral iSC.

We also analyzed the onset of the decrease in fixation-related activity during the gap interval across FEF inactivation. Like the result obtained for PREP activity, we found no change in the onset time of FIX activity across our sample of neurons (n=17; p=0.62; z-val=1.3416; Wilcoxon sign rank test).

### FEF inactivation produces a diversity of effects on the trial-by-trial relationship between PREP activity and the subsequent SRT

We now return to the observation that FEF inactivation reduces PREP activity in the caudal iSC, and examine how this relates to the eventual SRT. In particular, we are interested in whether FEF inactivation alters the trial-by-trial relationship between PREP and SRT, and whether the level of PREP activity attained prior to a SRT-matched saccade increases, decreases, or stays the same. Addressing these issues will shed light on the nature of how preparatory activity in the iSC relates to saccade initiation during FEF inactivation. We address these issues first at a neuron-by-neuron level, and then conduct a novel population-based analysis.

Consistent with previous studies (Dorris et al. 1997; Rezvani and Corneil 2008), prior to FEF inactivation most neurons exhibited a significantly negative trial-by-trial correlation between the final level of PREP activity and SRT, meaning that greater levels of PREP activity preceded the generation of shorter-SRT saccades. We term this the PREP-SRT relationship. This negative correlation reached significance in 21 of 35 (60%) of neurons before FEF inactivation. Examples of the PREP-SRT relationship are shown in Figs. 4A-C (red dots), showing data from the same neurons as shown in Fig. 2. Figure 4A&C show example of neurons where the PREP-SRT relationship was significant before FEF inactivation; Fig. 4B shows an example of a neuron that did not exhibit this relationship when the FEF was not inactivated. Across our sample, the r^2^ value of this relationship was 0.195± 0.21 (range 0 to 0.8165, see upper histograms in Fig. 4D), similar to previously published results.

**Figure 4:**
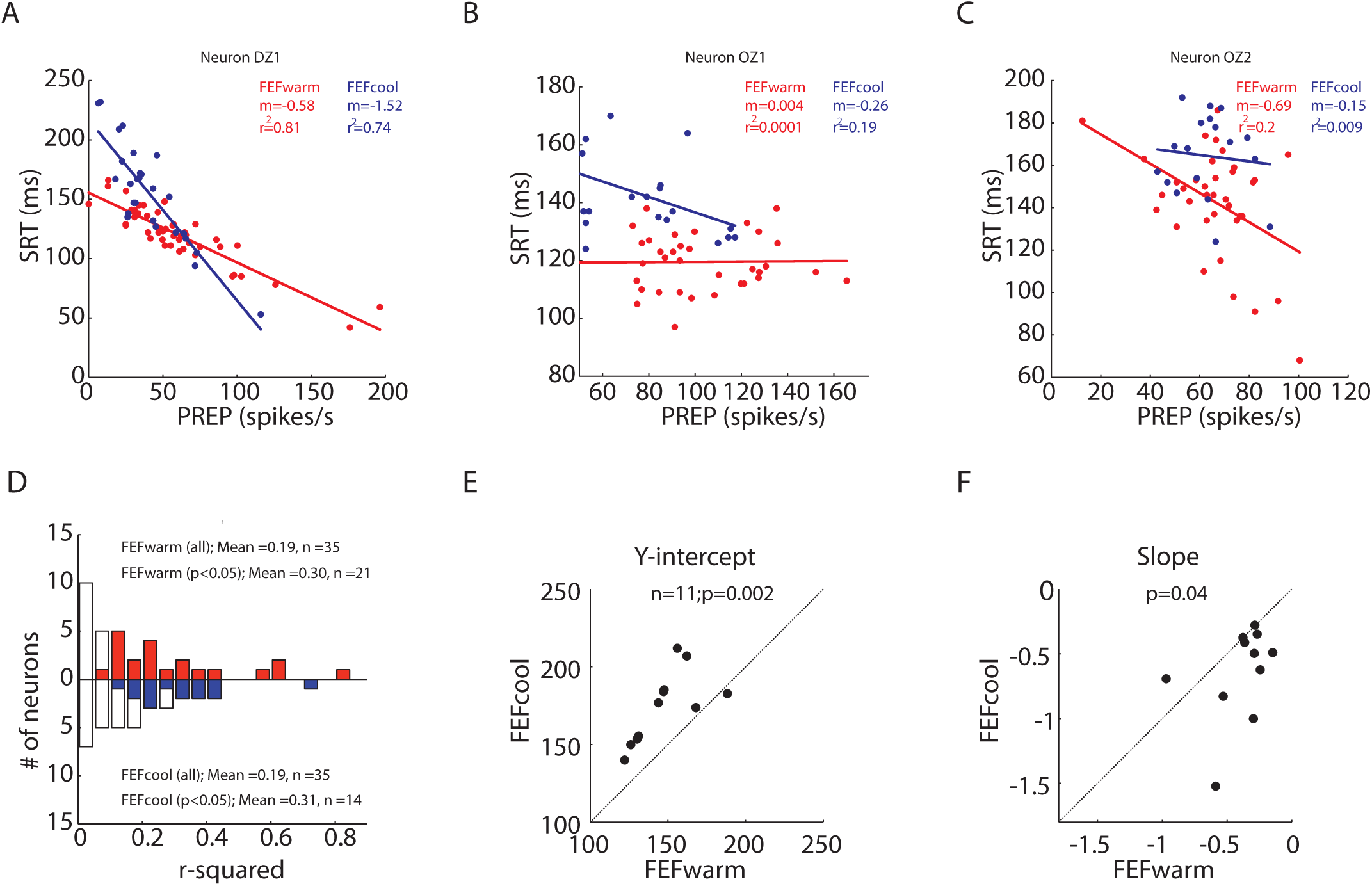
FEF inactivation changes trial by trial relationship between preparatory activity and SRT. (A, B & C) Trial by trial linear regression fits (solid lines) between PREP and SRT during FEF*warm* (red) and FEF*cool* (blue) for three example neurons shown in Fig.2. Each dot shows data from a single trial. (A) shows an increase in y-intercept and steepening of the relationship during FEF inactivation. (B) shows emergence of a PREP-SRT relationship during FEF inactivation. (C) shows the loss of a PREP-SRT correlation during FEF inactivation. (D) shows the changes in the coefficient of determination (r^2^) during FEF inactivation. Upright histogram shows FEF*warm* condition and inverted histograms shows FEF*cool* condition. Red and blue histograms indicate the r^2^ values for neurons that retained a significant PREP-SRT correlation during FEF*warm* and FEF*cool* conditions, respectively. (E & F) shows the changes in y-intercept (E) and slope (F) for the 11 neurons that exhibited significant PREP-SRT correlation during both FEF*warm* and FEF*cool* conditions.

How then did the relationship between PREP activity and the subsequent SRT change during FEF inactivation, particularly considering the overall reduction in the level of PREP activity? We are particularly interested in how many neurons retained a significant PREP-SRT relationship, whether there was any change in the amount of SRT variance explained by PREP variance, and whether there were any changes in the parameters of a linear fit through the PREP-SRT relationship (e.g., slope, y-intercept).

Surprisingly, FEF inactivation produced a diversity of effects on the PREP-SRT relationship, with the relationship either remaining (Fig. 4A, blue lines), emerging (Fig. 4B), or disappearing (Fig. 4C). The example in Fig. 4C is interesting, because FEF inactivation did not change PREP activity in this neuron (Fig. 2C), but did abolish the PREP-SRT relationship. Across our sample, 14 of 35 (40%) neurons exhibited a significant PREP-SRT correlation during FEF inactivation, representing decrease of 20% compared to the percentage of significant neurons during the FEF warm condition. Of these 14 neurons, 11 exhibited a significant PREP-SRT relationship in the FEF warm and FEF cool conditions (e.g., like the neuron in Fig. 4A), whereas 3 other neurons exhibited the relationship only in the FEF cool condition (e.g., like the neuron in Fig. 4B).

Across the sample of iSC neurons, the amount of variance in SRT explained by final level of the PREP activity (e.g., the r^2^ value of the PREP-SRT correlation) did not differ across the FEF warm versus cool conditions, regardless of whether we considered all 35 neurons (e.g., comparing the upward and downward histograms in Fig. 4D; p=0.63, z-val=-0.475; Wilcoxon sign sum test), or only the subset of neurons exhibiting a significant PREP-SRT correlation on an individual neuron basis (e.g., the colored histograms in Fig. 4D; p=0.49, z-val=-0.6903). For the 11 neurons that exhibited a significant correlation in both the FEF warm and FEF cool conditions, we found significant increases in both the y-intercept (Fig. 4E, p=0.002; z-val=-2.8451; Wilcoxon sign rank test) and a steepening of the slope (Fig. 4F, p=0.04; z-val=2.0449). Effectively, these changes mean that the linear regressions were pivoting around the lower SRT ranges, so that progressively greater levels of PREP activity preceded longer-SRT saccades.

Taken together, the effect of FEF inactivation on the PREP-SRT relationship was somewhat inconsistent: although this relationship was maintained in a subset of neurons, it was either gained or lost during FEF inactivation in others. Given this neuron-by-neuron diversity, it is difficult to draw any conclusions in how PREP activity in the caudal iSC relates to saccade initiation when the FEF is inactivated. We therefore conducted the following population-based analysis, to specifically examine iSC activity in relation to saccades generated at similar SRTs.

### Across the population, the final level of iSC preparatory activity is unaltered during FEF inactivation of SRT-matched saccades

The logic of our population-based analysis is shown in Fig. 5. This analysis, which exploits the overlap in SRT distributions during the FEF warm and FEF cool conditions, was conducted separately for each monkey. First, we constructed the SRT distributions for the FEF warm and FEF cool conditions (red and blue histograms respectively, in Figs 5A&C), pooling SRTs across all contraversive trials in which PREP activity was also recorded. The increases in SRTs during FEF inactivation is shown by the rightward shift of the blue compared to red histograms (Monkey DZ: p=8.18e-046; z-val=14.2079; Monkey OZ:p=8.84e-009; z-val=5.7516; Wilcoxon ranksum test). We then identified a 50ms range where the SRT distribution in the FEFwarm and FEF cool conditions overlapped, with at least 5% of the total trials within each 10 ms bin. The range of overlapping SRTs was 110 to 160ms for monkey DZ (shaded region, Fig. 5A) and 120 to 170ms for monkey OZ (shaded region, Fig. 5C). After identifying the appropriate range, we then subdivided the SRTs into partially overlapping sub-populations, constructing 7 subpopulations of 20ms bins incrementing by 5ms (e.g., 110-130ms, 115-135ms,…until 140-160ms for monkey DZ). Using the trials corresponding to the individual SRTs within each bin, we pooled iSC activity across all neurons, and then constructed population averages. For example, the spike density functions shown for the 110-130ms bin for monkey DZ (leftmost functions in Fig. 5B) were constructed from activity recorded from all iSC neurons on trials where the SRT fell into this range. This analysis allows us to construct comparable snapshots of iSC PREP activity across the population attained before various ranges of SRTs in either the FEF warm or FEF cool conditions.

**Figure 5:**
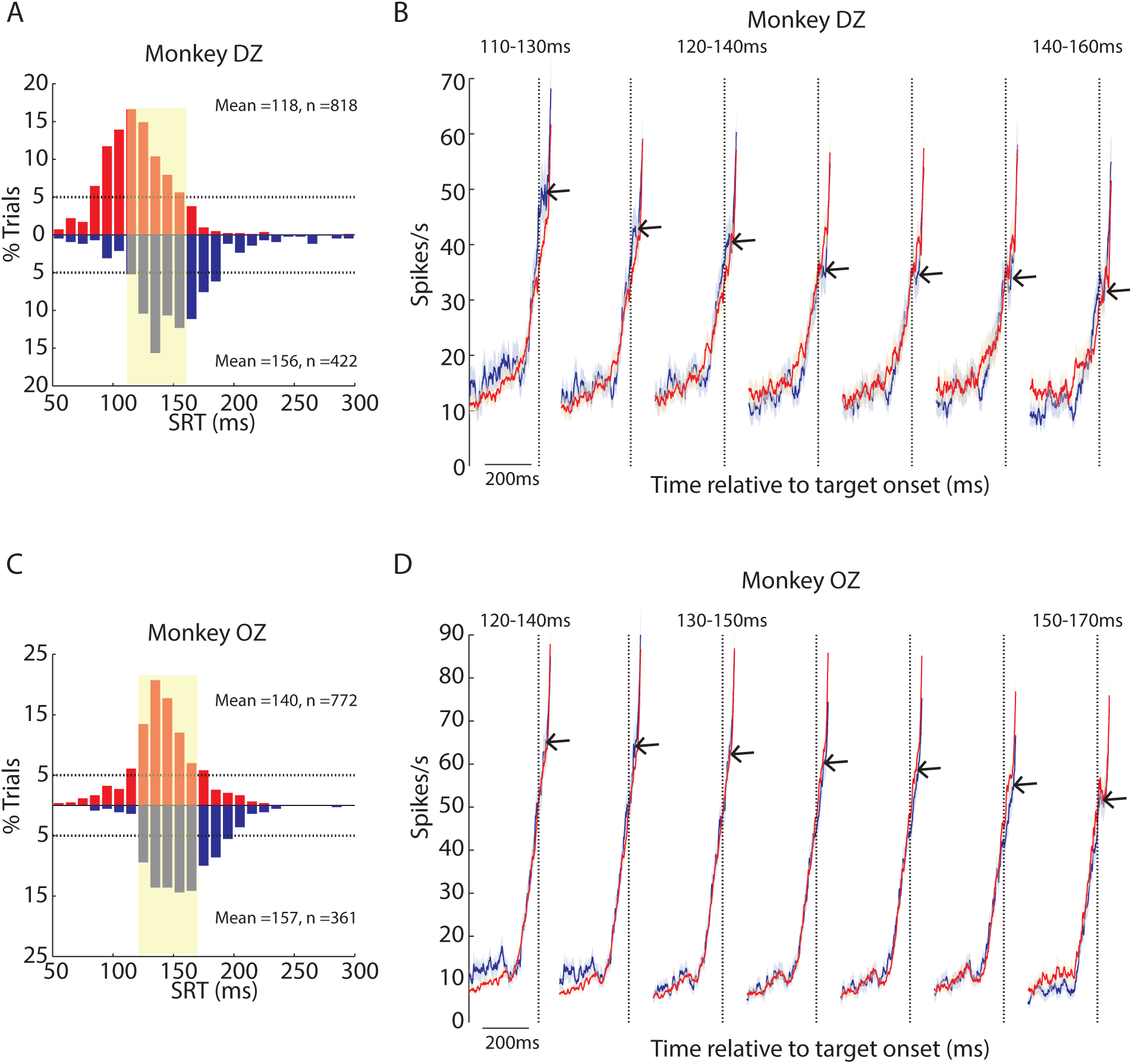
FEF inactivation did not alter population preparatory activity in the iSC for SRT matched saccades. (A & C) shows the SRT histograms for FEF*warm* (red and upright) and FEF*cool* (blue and inverted) conditions for monkey DZ and monkey OZ, respectively. The shaded region indicates the 50ms range of SRT in both monkeys where enough trials (> 5% trials in each 10ms SRT bins; the horizontal black dotted lines) contributed to both FEF*warm* and FEF*cool* conditions. (B & D) shows the profiles (Mean ± SE) of population iSC preparatory activity at different SRT ranges for monkey DZ and monkey OZ, respectively. Population profile for FEF*warm* (red) and FEF*cool* (blue) condition overlapped across the range of SRT studied.

The population spike density functions arising from this analysis are shown in Figs. 5B&D for monkey DZ and monkey OZ, respectively. Data is aligned on target onset (dotted vertical line), and is zoomed in on the time epoch encompassing the baseline and PREP intervals (e.g., visual and saccade-related activity not shown). We emphasize many features from this analysis. First, regardless of whether the FEF was inactivated or not, the final level of PREP activity, extracted from 10 to 30ms after target onset, decreased with increasing SRTs (for example see arrows pointing to final PREP activity during FEF inactivation (blue profiles) in Figs. 5B&D); this is another way of representing the inverse relationship between PREP and SRT. Second, when matched for SRT, the population iSC PREP activity appears largely unaltered during FEF inactivation: note from Fig. 5B (monkey DZ) and Fig. 5D (monkey OZ) how the blue and red activity profiles overlap almost perfectly for all the SRT bins. To quantify these observations, we extracted the baseline, slope of PREP accumulation, and final PREP level of activity as was done for the single-neuron analysis shown in Fig. 2. Doing so revealed no significant change in any parameter for any range of SRT during FEF inactivation (p > 0.05 for all comparisons even when not correcting for multiple comparisons, Wilcoxon rank sum test).

In summary, this population level analysis revealed that the overall level of iSC preparatory activity was unaltered during FEF inactivation, providing one compares saccades matched for SRT.

### Rare but precious: preserved iSC activity during express saccades

The SRT ranges studied in the above population analysis did not include express saccades, which we defined as SRTs between 70-120ms in accordance to previous studies (Schiller et al. 1987; Paré and Munoz 1996; Dorris et al. 1997; Sparks et al. 2000). As shown by the SRT distributions in Fig. 5A & C, both monkeys still generated express saccades during FEF inactivation, although their incidence was reduced. To analyze such rare saccades, we adopted the following SRT-matching logic. First, we identified those rare trials where a contraversive express saccade was generated during FEF inactivation. Using the SRT of this “FEF cool” express saccade, we then searched the FEF warm data recorded from the same iSC neuron for trials where a matching “FEF warm” contraversive express saccade was generated with a SRT within ± 3ms. Across our entire sample, we obtained 191 such matches (165 from monkey DZ and 26 from monkey OZ). We then pooled iSC activity across all matches to produce population-level representations of PREP activity preceding express saccades generated with the FEF inactivated or not. As shown in Fig. 6A, FEF inactivation had no effect on the final level of iSC PREP activity (Fig. 6D; p=0.59; z-val=-0.5287; Wilcoxon sign rank test). However, the level of baseline was significantly increased during FEF inactivation (Fig. 6B; p=0.0014; z-val=-3.1871; Wilcoxon sign rank test) and the slope of PREP accumulation also showed an increase (Fig. 6C; p=0.04; z-val=-2.0317; Wilcoxon sign rank test. This increase in baseline activity resembles that observed for the lowest SRT bins in the preceding analysis (i.e., compare baseline activity across inactivation in the left-most spike density functions in Figs. 5B&D), although a comparison of baseline activity for the lowest SRT range did not reach significance in either animal.

**Figure 6:**
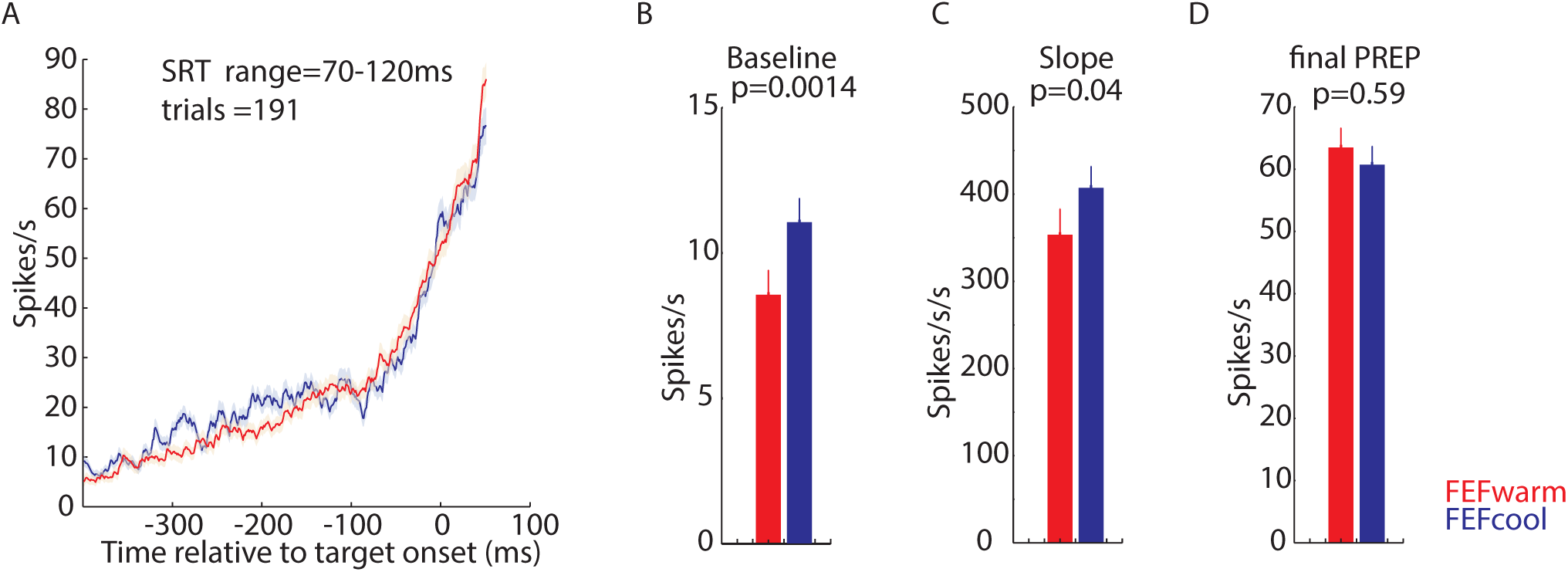
FEF inactivation did not alter the final level of iSC population PREP activity for latency matched express saccades. (A) shows the iSC population profiles for express saccades with matched pairs of SRT within ± 3ms for FEF*warm* (red) and FEF*cool* (blue) conditions. Across our sample, 191 matched trials could be extracted from both monkeys. (B, C & D) shows a significant increase in baseline activity (B) and slope of PREP activity (C) but an unaltered final level of PREP activity (D) (Wilcoxon sign rank test).

In summary, express saccades during FEF inactivation, although reduced in overall incidence, were associated with higher baseline activity but an unchanged final level of activity at the end of PREP epoch.

## Discussion

We combined unilateral cryogenic inactivation of the FEF with recordings of low frequency activity in the ipsilesional iSC, to better understand the coordination of these structures during saccade preparation, and the way in which iSC preparatory activity relates to saccade initiation when the FEF is compromised. As expected, FEF inactivation increased contralateral SRTs (Keating and Gooley 1988a; Sommer and Tehovnik 1997; Dias and Segraves 1999; Peel et al. 2014), and decreased the overall level of iSC preparatory activity, consistent with a loss of excitatory inputs into the caudal iSC. At an individual neuron level, we observed a diversity of effects of FEF inactivation on the inverse relationship between iSC preparatory activity and SRT, with this relationship being characterized by an increased intercept and steepness (slope) in some neurons and abolished in others. However, the collective population output of iSC preparatory activity was unaltered for SRT matched saccades, including those in the range of express saccades. Complementing these findings, fixation-related activity in the rostral iSC generally decreased during FEF inactivation, consistent with a loss of excitatory input into the rostral iSC. Here, we consider the implications of these results in terms of the functional contribution of the FEF during saccade preparation, and the pathways by which such a contribution could be realized.

### Reduction in preparatory iSC activity during FEF inactivation relates to a withdrawal of excitatory inputs rather than a rebalance of iSC activity

There are many potential mechanisms by which large-volume FEF inactivation could alter the level of iSC preparatory activity and increase contralesional SRTs. Given what is known about the anatomy and functional content of signals relayed directly from the FEF to the iSC (Künzle et al. 1976; Segraves and Goldberg 1987; Keating and Gooley 1988a; Stanton et al. 1988; Sommer and Tehovnik 1997; Tian and Lynch 1997; Dias and Segraves 1999; Everling and Munoz 2000; Sommer and Wurtz 2001), inactivation of a region within the FEF may produce a topographically-aligned withdrawal of excitatory inputs into the corresponding region of the iSC. This explanation may also explain the reduction in fixation-related activity in the rostral iSC, providing our cooling loops inactivated the lateral region of the FEF associated with visual fixation (Izawa et al. 2004, 2009). Simultaneously, FEF inactivation may alter signaling in interconnected cortical (e.g. dorsolateral prefrontal cortex (DLPFC), supplementary eye fields (SEF), or lateral intraparietal area (LIP)) or subcortical (e.g., basal ganglia) inputs to the iSC, with the net effect of decreasing excitatory inputs to the iSC. Regardless of the exact mechanism, our core findings of reduced preparatory- and fixation-related activity in the caudal and rostral iSC, respectively, relates to our recent work showing that FEF inactivation reduces visual-, delay-period, and saccade-related activity (Peel et al. 2017). This recent work also correlated increases in SRT during FEF inactivation with delays in the onset of saccade-related accumulation during a delayed-saccade task. In the current study we did not find any changes in the onset of preparatory activity during FEF inactivation. As outlined below, our data suggests that SRT in the gap saccade task is largely dictated by the magnitude of preparatory activity attained just before arrival of the visual transient, rather than the time at which such activity starts to accumulate.

Could this generalized reduction in iSC activity relate to a redistribution of activity toward the contralesional iSC? Several observations argue against this interpretation. First, ipsilesional SRTs do not decrease during large-volume cryogenic inactivation or ablation of the FEF (Fig. 1A,C; see also (Peel et al. 2014; Kunimatsu et al. 2015)), in contrast to the “push-pull” rebalancing of oculomotor activity during small-volume pharmacological FEF inactivation (Sommer and Tehovnik 1997; Dias and Segraves 1999), Further, our recent study showed that contralesional iSC activity did not increase in delayed-saccade tasks during unilateral FEF inactivation (Peel et al. 2017). We also observed that rostral iSC activity decreases, rather than increases, upon unilateral FEF inactivation, suggesting that the decrease in caudal iSC preparatory activity is also not a consequence of rostro-caudal reciprocal inhibitory interactions. These observations from rostral iSC recordings are consistent with our previous observations of how the peak velocities of microsaccades decrease, rather than increase, during FEF inactivation (Peel et al. 2016).

### The level of preparatory activity reached by the end of the gap interval is unchanged during FEF inactivation for SRT-matched saccades

We also addressed, for the first time, if and how the negative relationship between iSC preparatory activity and SRT is influenced when a major input to iSC is suddenly removed. At the single neuron level, the effects of FEF inactivation were surprisingly diverse, with the relationship being abolished in some neurons and systematically altered (increased slope and y-intercept) in others. We recently reported a similar diversity of effects of FEF inactivation on saccade-related activity (Peel et al. 2017). We have no ready explanation for this diversity of effects. It may be that the FEF only projects directly to a subset of iSC neurons; however, to our knowledge there is no anatomical data that speaks directly to this point.

Despite differences in the effects of FEF inactivation on the single neuron, we found that the level of iSC preparatory activity across the population was unaltered during FEF inactivation, providing one compares SRT-matched saccades. Our interpretation is that while FEF is one important source of top-down excitatory input to iSC, it is not the sole source. Presumably, when inputs from other cortical areas like the DLPFC, SEF and LIP (Yamamoto et al. 2004; Koval et al. 2011; Chen et al. 2013; Johnston et al. 2014) and subcortical areas like basal ganglia and cerebellum (Wurtz and Hikosaka 1986; Hikosaka et al. 1989; Ashmore and Sommer 2013; Ohmae et al. 2017) bring iSC preparatory activity to a particular level, this level is associated with a given SRT, irrespective of the presence or absence of the FEF. This may be because, in the gap saccade task studied here, SRT is largely dictated by the merging of the visual transient with pre-existing preparatory activity (Dorris et al. 2007). Consistent with this, the effects of FEF inactivation on the visual transient recorded in the iSC during a delayed saccade task is relatively modest, with no changes in the timing and only small decreases in magnitude (Peel et al. 2017). Further, parametric manipulations of either the level of preparatory activity (Dorris et al. 1997; Basso and Wurtz 1998; Rezvani and Corneil 2008), the timing of vigor of the visual transient (Bell et al. 2006; Li and Basso 2008; Marino et al. 2012), or the interaction between these two signals (Dorris et al. 2007) all systematically alter SRT.

Our observation of an unaltered population profile for preparatory signals also extends to express saccades. Behavioral (Fischer et al. 1984; Paré and Munoz 1996) and neurophysiological (Edelman and Keller 1996; Dorris et al. 1997; Sparks et al. 2000) studies have emphasized a paradoxical nature of these movements: on one hand, they are the oculomotor expression of a low-level visual grasp reflex, but on the other hand their prevalence hinges on advanced saccadic preparation mediated by top-down inputs. FEF inactivation reduced, but did not abolish, the prevalence of express saccades, indicating that the FEF is an important, but not critical nor exclusive, source of top-down input favoring express saccade production. This observation complements lesion work in monkeys (Schiller et al. 1987) and clinical work in stroke patients (Rivaud et al. 1994) showing a persistent ability to generate express saccades despite permanent damage to the FEF, and shows that such persistence does not arise simply due to long-term compensation.

### A negligible role for the projection of the FEF to the saccadic brainstem during saccade preparation

An unaltered level of population preparatory activity also allows us to speculate on the nature of FEF projections that bypasses the iSC and directly contacts the oculomotor brainstem. Although, the existence of this *bypass pathway* from FEF to brainstem burst generator is supported by anatomical and antidromic studies (Leichnetz 1980, 1981; Huerta et al. 1986; Segraves 1992), whether this pathway has any substantial role in saccade generation in the intact animal is not clear. The presence of this bypass pathway is thought to aid recovery of oculomotor functions following iSC ablations (Schiller 1977; Keating and Gooley 1988b). If signals carried along the bypass pathway would have had a permissive role in saccade initiation, as suggested by the antidromic work of Segraves (1992), then loss of such signaling during FEF inactivation would have required a higher-level of iSC preparatory activity for a given SRT. Instead, since the level of iSC preparatory activity associated with a given SRT remains unchanged during FEF inactivation, our interpretation is that the bypass pathway plays a negligible role in saccade preparation in the intact animal. This interpretation supports conclusions in a study by Hanes and Wurtz (2001), showing that saccades evoked by FEF stimulation were abolished when the corresponding topographic location in the iSC was pharmacologically inactivated (Hanes and Wurtz 2001), suggesting that signals relayed via the bypass pathway were insufficient to initiate a saccade in an intact animal.

## Acknowledgements

This work was supported by operating grants from the Canadian Institutes of Health Research to BDC (MOPs: 93796, 123247 and 142317) and the Natural Sciences and Engineering Research Council (NSERC; RGPIN-311680). SD and TRP were supported by funding from an NSERC CREATE grant and TRP was supported by an Ontario Graduate Scholarship.

